# Retinal cell mosaics in the valproate-induced rat model of autism spectrum disorder

**DOI:** 10.64898/2026.06.14.732149

**Authors:** Ildikó Telkes, Katalin Fusz, Tibor Zoltán Jánosi, Péter Kóbor, Abdurrahman ElZafarany, Zoltán Sári, Kristóf László, Péter Buzás

**Author notes:** **Corresponding author:** Péter Buzás, Institute of Physiology, Medical School, University of Pécs, Szigeti út 12., 7624 Pécs, Hungary.

## Abstract

Valproic acid (VPA) is a widely used antiepileptic drug that also increases the risk of neurodevelopmental disorders in the offspring of exposed mothers. Prenatal exposure to VPA is a widely used rodent model of autism spectrum disorder (ASD). Anatomical, functional and molecular alterations in the retinas of various ASD model animals have been described, but the impact on the neural composition of the retina remains unclear. We examined whether the density and spatial regularity of selected retinal neuron populations are altered in the VPA induced model of ASD. Whole-mount retinas of 2-month-old VPA-treated and control animals were immunolabeled for S-cones, horizontal cells, AII amacrine cells, and parvalbumin-positive wide-field amacrine cells (PV-wfACs), and the positions of labelled cells mapped in various regions of interest across the retina. Multivariate analysis of the densities of the four cell types was used to identify latent factors accounting for their covariance structure. The first principal component was identified as a measure of retinal centrality and used to adjust for retinal location when testing for treatment effects. Linear mixed-effects modeling revealed a 9% decrease in AII amacrine cell density and a 12% increase in PV-wfACs density in VPA-model animals. Regularity indices calculated from nearest neighbor distances or Voronoi-domain areas of cell mosaics were largely unchanged. These findings suggest that prenatal VPA exposure selectively alters the cellular composition of inhibitory inner retinal circuitry in the rat ASD model, while leaving the self-organizing mechanisms responsible for retinal spatial order largely intact.

**Lay Summary:** Valproic acid (VPA) is a medicine for epilepsy, but it can also raise the risk of autism in children when taken during pregnancy. In rats exposed to VPA before birth, we found changes in certain nerve cells of the retina: one type of cell important for night vision was reduced, while another type increased slightly, while most other cells stayed the same. This suggests that the changes in development that lead to autism may also be reflected in the structure and function of the eye.

## Introduction

The ability of the retina to process images relies on its photoreceptors and various neuron types sampling the image at distinct spatial resolutions. Each cell type creates its own cellular mosaic, characterized by cell density and regularity. While average cell density is a large-scale measure that varies smoothly across the retina (such as between central and peripheral regions), regularity pertains to the local variation in the distance between neurons.

Retinal cell mosaics develop on the basis of genetic mechanisms controlling morphogenetic factors, cell migration and apoptosis (Sharma and Johnson 2000; Jusuf et al. 2012; Amini et al. 2018; Kozlowski et al. 2024) as well as self-regulating lateral interactions between neurons (Reese and Keeley 2016). Thus, perturbations of these mechanisms can potentially affect the density and regularity of retinal mosaics (Eglen and Willshaw 2002; Raven et al. 2003). Mutations of several genetic loci have been shown to alter either neuron density or the regularity of cell mosaics in various, often highly specific ways (Williams et al. 1998; Strettoi and Volpini 2002; Kay et al. 2012; Jusuf et al. 2012; Kozlowski et al. 2024). Perturbations of neural development caused by teratogens or epigenetic factors can also affect the adult retina, although the changes have been found to be less specific, and often involve other ocular structures (Tandon and Mulvihill 2009).

Valproic acid (VPA) is a widely used antiepileptic, with additional indications for bipolar disorder and migraine (Tursunov et al. 2023). Its anticonvulsive property is largely explained by its increasing gamma amino butyric acid (GABA) neurotransmission through various mechanisms (Tursunov et al. 2023). However, VPA is also teratogenic, increasing the risk of major congenital malformations (such as neural tube defects and cardiac anomalies (Nau 1994)), decreased intelligence quotient and neurodevelopmental disorders including autism spectrum disorder (ASD) in a dose-dependent manner (Ornoy 2009) in the children of exposed mothers. The teratogenicity of VPA appears to derive to a large extent from its histone-deacetylase inhibitor activity, which results in changes in gene expression, although direct perturbations of neurotransmitter systems have also been implicated (Silvestrin et al. 2013).

Autism spectrum disorder (ASD) is a multifactorial neurodevelopmental condition characterized by impairments in social interaction and repetitive behaviors (American Psychiatric Association 2013). A recent meta-analysis estimated that the median prevalence of autism is 100/10 000 globally, with a median male/female ratio of 4.2 (Zeidan et al. 2022). The pathomechanism of ASD is, with certainty, multifactorial but a common final pathway still remains unknown. A wide range of biological, environmental, and genetic factors have been identified that increase susceptibility to ASD (Bailey et al. 1995; Lord et al. 2018; Kavitha et al. 2021). Among various animal models, prenatal VPA treatment is widely used due to its ability to reliably induce ASD-like traits, particularly in male offspring (Nicolini and Fahnestock 2018).

In rodents, VPA treatment given around embryonic days 12-13 maximizes the likelihood of ASD-like symptoms in the offspring. This developmental time window also coincides with the first wave of retinal neurogenesis (Rapaport et al. 2004). In a recent study (Guimarães-Souza et al. 2019), functional and molecular alterations of the retinas of VPA-induced ASD model mice have been described. These experiments suggested impaired photoreceptor function, as evidenced by a decreased a-wave of the electroretinogram relative to controls. Furthermore, the expression of molecular markers associated with GABAergic neurotransmission was also reduced.

Here, we were interested in whether the mosaics of certain retinal neurons are altered in adult rats following prenatal VPA exposure. We chose horizontal cells, parvalbumin-positive amacrine cells including AII, and S-cones for this analysis to probe different retinal processing systems and different levels of circuitry. Horizontal cells and amacrine cells are the main interneurons of the outer and inner plexiform layers, respectively, forming hubs at successive levels of the retinal circuitry. AII amacrine cells specifically, are the most frequent amacrine cell type, which contribute to the processing of rod-as well as cone signals (Demb and Singer 2012; Strettoi et al. 2018). The motivation for including S-cones in the set was two-fold: 1. Since all cones and rods are thought to differentiate from S-opsin expressing precursors, a perturbation of early neural development might be reflected in the adult S-cone mosaic (Szél et al. 1994; Lukáts et al. 2005; Hussey et al. 2022). 2. S-cone signals are processed in specialized retinal pathways largely independent of most other circuits, which are M/L-cone-dominated (Calkins and Sterling 1999; Miyagishima et al. 2014). Furthermore, to detect treatment-induced alterations in multiple retinal cell populations despite substantial topographic variability, we developed a statistical framework for identifying and comparing equivalent locations across retinas.

## Materials and Methods

### Animals and sample preparation

To create the autism model, pregnant Wistar rats (*Rattus norvegicus*, Table 1) were treated with 500 mg/kg of body weight valproic acid (VPA) on day 12.5 of pregnancy. Autism-like traits were verified with a three-compartment social interaction test and anxiety was verified with an elevated-floor cross maze test (László et al. 2022a, b). In the three-compartment social interaction test, VPA-treated rats spent significantly less time in social interaction than control animals (VPA: 25.67 ± 2.67 s, n = 6; control: 84.75 ± 7.04 s, n = 4; t(3.88) = 7.85, p<0.05 in t-test). In addition, VPA-treated animals spent significantly less time in the open arms of the elevated plus maze (VPA: 12.54 ± 1.79 s; control: 36.22 ± 3.31 s; t(8) = 6.89, p<0.001 in t-test), indicating increased anxiety-like behavior. These findings confirm the presence of a robust behavioral phenotype in the VPA-treated animals used in the present study.

**Table 1.**
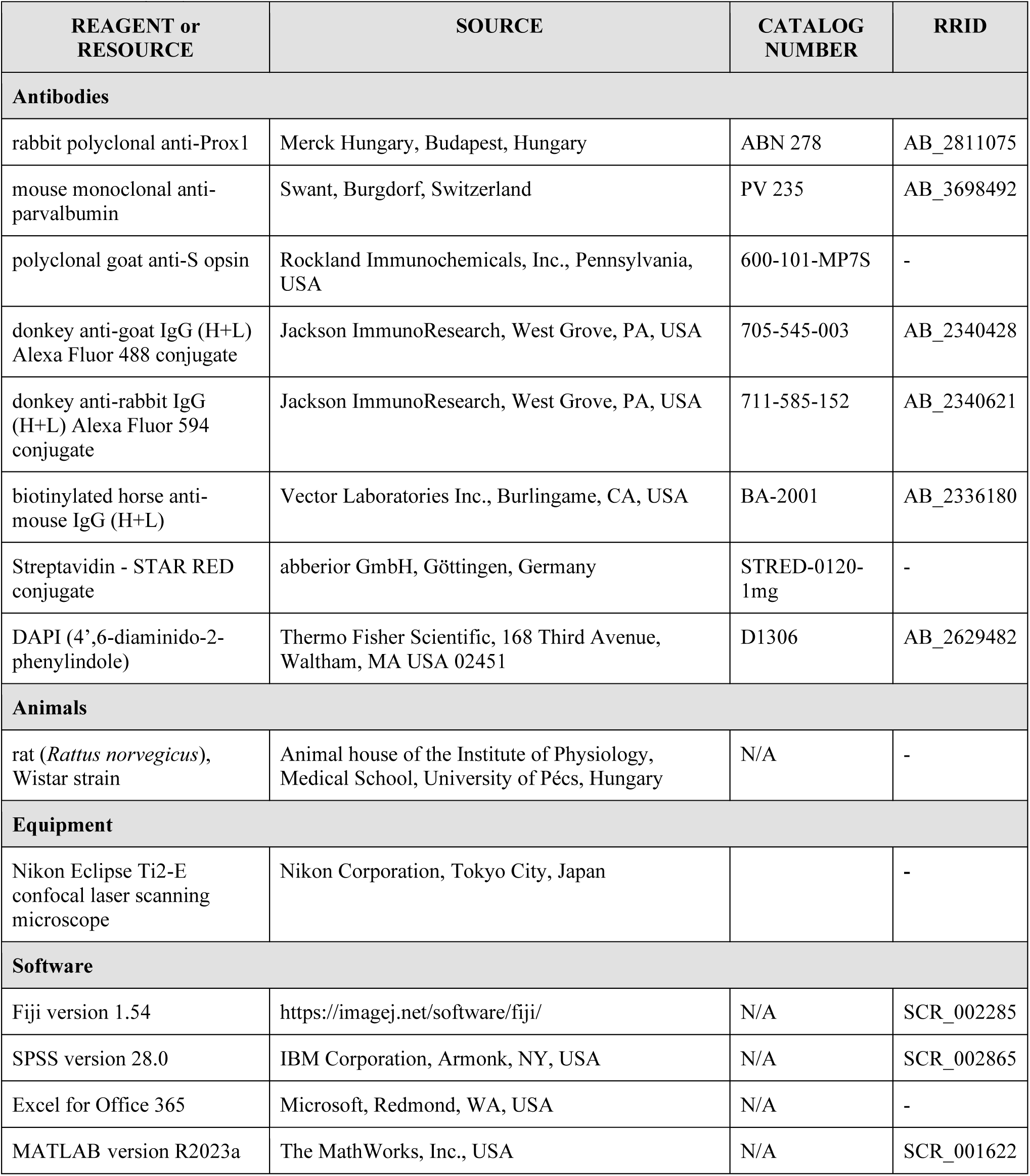
List of key resources.

| REAGENT or RESOURCE | SOURCE | CATALOG NUMBER | RRID |
| --- | --- | --- | --- |
| **Antibodies** |  |  |  |
| rabbit polyclonal anti-Prox1 | Merck Hungary, Budapest, Hungary | ABN 278 | AB_2811075 |
| mouse monoclonal anti-parvalbumin | Swant, Burgdorf, Switzerland | PV 235 | AB_3698492 |
| polyclonal goat anti-S opsin | Rockland Immunochemicals, Inc., Pennsylvania, USA | 600-101-MP7S | - |
| donkey anti-goat IgG (H+L) Alexa Fluor 488 conjugate | Jackson ImmunoResearch, West Grove, PA, USA | 705-545-003 | AB_2340428 |
| donkey anti-rabbit IgG (H+L) Alexa Fluor 594 conjugate | Jackson ImmunoResearch, West Grove, PA, USA | 711-585-152 | AB_2340621 |
| biotinylated horse anti-mouse IgG (H+L) | Vector Laboratories Inc., Burlingame, CA, USA | BA-2001 | AB_2336180 |
| Streptavidin - STAR RED conjugate | abberior GmbH, Göttingen, Germany | STRED-0120-1mg | - |
| DAPI (4',6-diaminido-2-phenylindole) | Thermo Fisher Scientific, 168 Third Avenue, Waltham, MA USA 02451 | D1306 | AB_2629482 |
| **Animals** |  |  |  |
| rat (*Rattus norvegicus*), Wistar strain | Animal house of the Institute of Physiology, Medical School, University of Pécs, Hungary | N/A | - |
| **Equipment** |  |  |  |
| Nikon Eclipse Ti2-E confocal laser scanning microscope | Nikon Corporation, Tokyo City, Japan |  | - |
| **Software** |  |  |  |
| Fiji version 1.54 | https://imagej.net/software/fiji/ | N/A | SCR_002285 |
| SPSS version 28.0 | IBM Corporation, Armonk, NY, USA | N/A | SCR_002865 |
| Excel for Office 365 | Microsoft, Redmond, WA, USA | N/A | - |
| MATLAB version R2023a | The MathWorks, Inc., USA | N/A | SCR_001622 |

The control group consisted of the offspring of untreated females because our goal was to characterize the established VPA-induced ASD model as it is commonly used, rather than to isolate the pure pharmacological effect of valproic acid. Animals assigned to the same experimental group were not littermates.

Altogether, the retinas of 10 male, two-months-old adult rats collected: six from the treated group and four from the control group. Rats were transcardially perfused with phosphate-buffered saline (PBS) and then with 4% paraformaldehyde following anesthesia with ether and subsequent lethal injection of T61 (embutramide 250 mg/kg, tetracaine HCl 6.25 mg/kg, mebezonium iodide 63 mg/kg, Intervet, Boxmeer, The Netherlands). The eyes were removed and cut along the *ora serrata.* The lens and vitreous body were eliminated. The posterior eyecups were postfixed overnight at +4 °C in 4% paraformaldehyde in PBS and the retinas were prepared in cold PBS.

Animals were housed (two per cage) in a temperature- and light-controlled room (22 ± 2 °C; 12:12 h light-dark cycle with lights on from 6:00 a.m.). Standard laboratory food pellets (CRLT/N standard rodent food pellet, Charles River, Budapest, Hungary) and tap water were available ad libitum. All animals were housed and experiments conducted in compliance with Hungarian and European legislation. All procedures were approved by the Directorate for Food Chain Safety and Animal Health of the Baranya County Government Office, Hungary (permit number BA02/2000-18/2023).

### Immunohistochemistry and confocal microscopy

Free-floating retinal quadrants or whole retinas were first incubated with blocking solution composed of 10% normal goat serum in antibody diluting solution (0.25% bovine serum albumin, 0.001% sodium azide, and 0.2% Triton X-100 in 0.1 M PBS) for 2 days. The same solution was used for all further antibodies unless stated otherwise. Tissue samples were then incubated with the primary antibodies at +4 °C for 4 days using the following dilutions: polyclonal anti-Prox1 produced in rabbit (C-terminal peptide of human PROX1), 1:2000; monoclonal anti-parvalbumin produced in mouse (antigen: parvalbumin purified from carp muscle), 1:6000; polyclonal anti-S opsin produced in goat (antigen: recombinant human blue opsin), 1:200. The labelling corresponded with the well-known patterns for these markers as described in the Results (Figure 1) and Discussion (Table 7).

**Figure 1.**
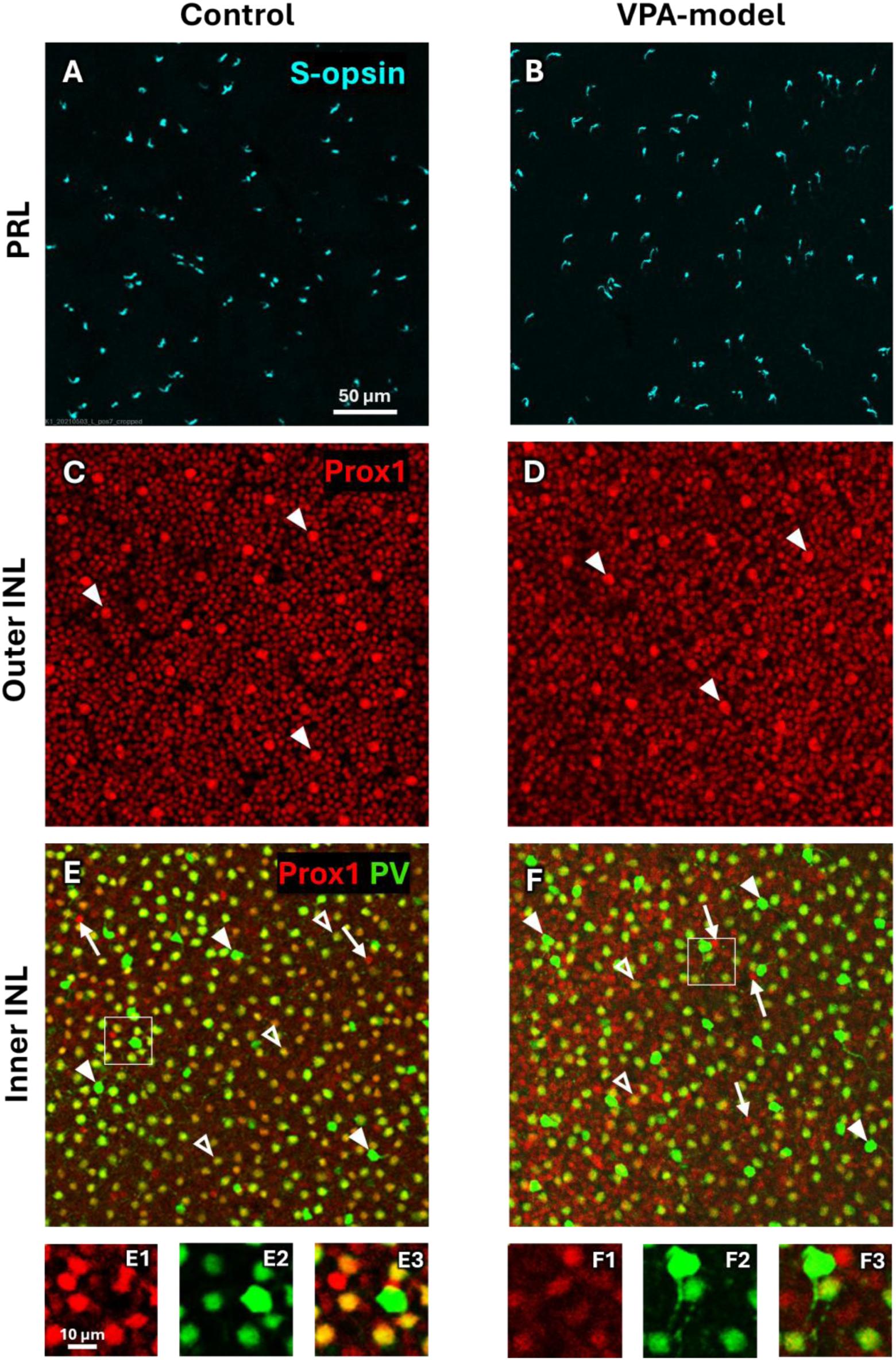
S-opsin, Prox1 and parvalbumin (PV) labelling in the retinas of control (left column) and VPA-induced autism model (right column) rats. Both regions of interest were taken from the dorsal retina, the distances from the optic nerve head were 0.8 mm and 1.3 mm for the control and VPA-model examples, respectively. (A-B) In the photoreceptor layer (PRL), blue (S-) cones were visualized using anti-S-opsin antibody. (C-D) On the outer aspect of the inner nuclear layer (outer INL), anti-Prox1 antibody labelled the horizontal cell mosaic (HC, arrowheads) and many small ON-bipolar cells. (E-F) The deeper INL showed both PV (green channel) and Prox1 (red channel) labelling (inner INL). The most numerous cell type was the double labelled AII amacrine cell (open arrowheads). Strongly PV-positive cell bodies belong to a population of wide-field amacrine cells (PV-wfAC, solid arrowheads). Occasionally, Prox1-positive amacrine cells lacked PV immunoreactivity (arrows). E1-3 and F1-3 show regions of E and F respectively (white boxes), at higher magnification with split Prox1 (1), PV (2) and merged (3) channels. The scale bar of 50 µm applies to images E-F, the scale bar of 10 μm applies to images E1-3 and F1-3.

The following steps were done at +4 °C overnight. Prox1 immunoreactivity was visualized with donkey anti-rabbit Alexa Fluor 594 (1:100 dilution), parvalbumin (PV) immunoreactivity was visualized with biotinylated anti-mouse antibody (1:100 dilution) overnight, and then streptavidin conjugated STAR RED (1:200 dilution) also overnight. S-opsin immunoreactivity was visualized with donkey anti-goat Alexa Fluor 488 (1:500 dilution). We used DAPI (4’,6-diamidino-2-phenylindole) at 5 μg/ml concentration for visualizing the cell nuclei. The sources of antibodies are listed in Table 1.

Retinal pieces were washed between the incubations five times for 10 min in 0.1 M PBS and then mounted in Aqua-PolyMount (Polysciences, Warrington, PA, USA) or VectaShield (Vector Labs., USA) media with the ganglion cell layer facing the coverslip.

We inspected the flat-mounted retinas using a Nikon Eclipse Ti2-E confocal laser scanning microscope (Nikon Corporation, Tokyo City, Japan) through a 20× objective lens, following a tile-scan of the entire retina at 10× for localization. We took confocal z-stacks at regions of interest (ROIs, see below) selected from various regions of the retina (Supplementary Figure S1). The horizontal size of the ROIs was 135×135 μm and the z-stacks spanned depth from the photoreceptor layer to the optic fibers. Alternatively, the depth of the retina was scanned in two parts whereby the outer nuclear layer was skipped to save time. Voxel size was 0.132 μm × 0.132 μm × 0.381 μm or 0.132 μm × 0.132 μm × 0.5 μm.

### Data processing

The necessary number of ROIs was estimated by power analysis in G*Power version 3.1.9.7 (Faul et al. 2009). The estimates were based on calbindin-positive horizontal cell, amacrine cell and ganglion cell density data of *Engrailed-2* knock-out and wild-type mice published by Zhang et al. (2019). Assuming an effect size of 0.9 (estimated from their Figure 5) and *α* = 0.05, 34 observations per group would be required to reach 95% statistical power in an independent sample t-test. Because the final analysis treated ROIs as repeated observations nested within animals, this t-test based calculation should be regarded as a planning estimate rather than a formal power analysis for the final statistical model.

For the present study, we analyzed 71 ROIs (32 from controls, 39 from VPA-treated animals) sampled from 10 retinas (control n=4, VPA n=6). The number of ROIs per animal was 5, 6, 10 and 11 for controls and 1, 3, 4, 6, 8 and 17 for the VPA group (median = 6, also see Supplementary Figure S1). In a few ROIs, certain cell types could not be reliably identified. These ROIs were excluded from principal components analysis, where complete data across all four cell types were required, but they were retained for analyses involving the available cell types (e.g., bivariate correlations).

Z-stack images were further processed using the Fiji image analysis software package (version 1.54, Schindelin et al., 2012). Four retinal cell types were identified in the ROIs: S-cones, horizontal cells (HCs), PV-positive wide field amacrine cells (PV-wfACs) and AII amacrine cells. The distinguishing features of these cell types are described in the Results section. Their positions were recorded as 2-dimensional coordinates in the image plane by marking the identified cell bodies manually using the Cell Counter plugin. Every ROI was assessed by a second investigator for areas where a cell type could not be reliably identified due to factors such as out-of-focus imaging or weak immunostaining, and these regions were manually masked. Altogether, 83579 marked cells were included in the final dataset. Areal density was calculated by dividing the number of identified cells by the unmasked area of the ROI.

Nearest neighbor distances and Voronoi domain areas were calculated by using custom-written scripts in MATLAB (version R2023a, MathWorks, Natick, MA, USA). For nearest neighbor analysis, cells that were closer to the boundary of the ROI or to the edge of a masked region than to any other cell were excluded from the calculation. For Voronoi tessellation, those domains that had one or more vertex outside the valid region of the ROI were ignored. The regularity of cell mosaics was quantified using two metrics calculated for each ROI separately: the nearest neighbor regularity index (NNRI), defined as the ratio of the mean nearest-neighbor distance to its standard deviation, and the Voronoi-domain regularity index (VDRI), defined as the ratio of the mean Voronoi-domain area to its standard deviation.

To minimize sampling bias, the researchers selecting the ROIs had no information about the anatomical orientation or exact topography of the given retina, although they were aware of the identity of the animal. The researchers labelling the individual cells within the ROIs were not aware of the identity of the animal or the location of the given ROI within the retina.

### Localizing ROIs

The anatomical locations of ROIs were estimated only to guide our sampling strategy, namely that we aimed at covering similar distances and anatomical directions in both experimental groups. Importantly, the ROI locations were not included in the main statistical model due to reasons detailed in the Results.

The ROI locations were estimated in two steps. First, the distance of each ROI from the optic disc was measured on calibrated low-power images. We avoid referring to the distance from the optic disc as “eccentricity”, because it is not relevant in judging the projection of retinal location into the visual field.

Second, the orientation of the retinal preparations was estimated from the distribution of S-cones. S-cones have a characteristic dorso-ventral gradient in the rat retina with the highest densities in ventral, the lowest densities in dorsal retina (Szél and Röhlich 1992; Ortín-Martínez et al. 2010). For each retina, we created a montage of confocal scans of the S-opsin label to determine the distribution of S-cones across the artefact-free area of the specimen. This was done either across the entire surface by using the Fiji plugin LABKIT (Arzt et al. 2022) or by counting the number of S-cones in six 500 μm ×500 μm sampling boxes equally spaced along a circle drawn around the optic disc at a mid-range distance. In both cases, the areal density was calculated and vectors were formed whose length was the S-cone density and the angle was the sample direction from the optic disc. The mean direction of these vectors was taken as the ventral anatomical direction. The location of each ROI was thus represented in a polar coordinate system centered to the optic disc and the zero direction pointing towards the nose. The above procedure could not be completed for five ROIs from three retinas (two controls and one treated) due to technical reasons. The estimated ROI locations in the retinal coordinate system are shown in Supplementary Figure S1.

### Statistical analysis

Statistical tests were performed in IBM SPSS (version 30.0, IBM Corporation, Armonk, NY, USA) and custom-written scripts in MATLAB. Normality of the raw data was tested using Shapiro-Wilk tests for each cell type and each of the following measures separately: cell density, nearest neighbor regularity index (NNRI) and Voronoi-domain regularity index (VDRI). The distributions of cell densities did not differ significantly from normal. For the regularity indices, significant departure from normality was found in all cases except for NNRIs of the two amacrine cell types and the VDRIs of PV-wfACs. Levene’s test (based on the means) was used to compare the variances of the treatment groups. Significant heteroscedasticity (p < 0.05) was detected for both density and regularity-index measures of AII amacrine cells. However, the subsequent mixed-effects analyses are generally robust against departures from normality and variance homogeneity of the raw data.

Two main types of statistical tools were used in our analysis: correlation analysis and linear mixed-effects modeling. Correlation analysis was used to explore the covariance structure of the dataset. Pearson’s correlation coefficients were used to evaluate relationships between cell densities. Differences in correlation structure between groups were tested using Fisher’s Z-test for equality of pairs of correlation coefficients and Jennrich’s test for equality of correlation matrices (Jennrich 1970; Vermorken 2008). Principal component analysis (PCA) was then employed to identify latent factors correlated with cell density distributions. The first principal component (PC1), interpreted as a proxy for retinal centrality, was used in subsequent analyses to control for location-dependent variation. For the pairwise correlations of regularity indices, Spearman’s correlation coefficients are reported. Because multiple ROIs originated from the same animal, the above correlations may reflect both within-animal and between-animal sources of variation.

Linear mixed-effects modeling (LME) was needed in the inferential analysis to account for multiple ROIs per animal when testing treatment effects on cell densities and regularity indices. In these statistical models, each ROI was entered as one observation, treatment was a fixed-effect factor and animal identity was included as a random effect contributing a random intercept on the dependent variable. The model equation for the outcome variable *Y* was

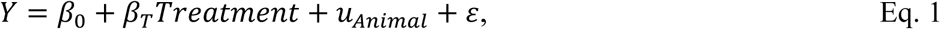

where *β*_0_ is the global intercept, *β_T_* is the treatment effect (VPA−Control), *u_Animal_* is the random intercept accounting for between-animal differences and *ε* is the residual error.

To adjust for the effects of location on cell density, a derived measure of retinal density gradient termed centrality (PC1, see below) was added in the model. Further, to separate the effects of between-animal and within-animal variation in centrality, PC1 was decomposed into animal-specific means (*PC1_mean_*, between-animal component) and deviations from those means (*PC1_cent_* = *PC1*−*PC1_mean_*, within-animal component), and both terms were entered as covariates into the linear mixed-effects model:

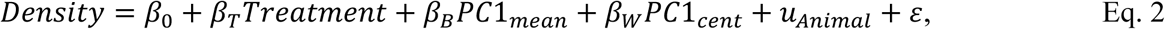

where *β_B_* and *β_W_* quantify the effects of between-animal and within-animal variation in centrality, respectively. Statistical interactions between treatment and the covariates were tested, but showed no significant contribution, therefore they were omitted from the model.

The goodness of fit of linear mixed-effects models was conveniently expressed by the estimated proportions of total variance explained by the model components. The variance proportion explained by the full model is quantified by the “conditional *R*^2^” (Nakagawa and Schielzeth 2013), which is the sum of variance proportions explained by the fixed effects and the random effect (animal identity). The former is denoted the “marginal *R^2^*” and the latter is denoted the “conditional intraclass correlation coefficient” (ICC) in the output of the MIXED procedure in SPSS (Anonymous 2022). Note that the conditional ICC is different from the conventionally used ICC, which is reported as “adjusted ICC” in SPSS. The adjusted ICC is defined as the proportion of the random effect variance within the sum of random effects and residual variance, and thus it does not take the variance explained by the fixed effects into account.

## Results

Figure 1 shows representative micrographs of two flat-mounted retinas triple-labelled for S-opsin, Prox1 and parvalbumin. The anti-S-opsin antibody revealed a sparse population of cones with strongly labelled outer segments (Figure 1 A, B). In the outer tier of the outer nuclear layer (ONL), S-opsin positive cell bodies appeared next to each outer segment (not shown). No other cell type was labelled by this antibody.

The inner nuclear layer (INL) was abundant with cell bodies labelled with the Prox1 or parvalbumin antibodies. The combination of these markers allowed the distinction of at least four cell types as described earlier (Wässle et al. 1993; Gábriel et al. 2004; Liu et al. 2022). Two types of cell nuclei expressing Prox1 could be readily distinguished on the outer aspect of the INL (Figure 1 C, D). Horizontal cells showed up as relatively sparse (mean cell densities 664±116 mm^−2^, n = 69 ROIs), larger in diameter and more intensely stained. Surrounding them, a dense array of bipolar cell nuclei could be distinguished.

The deeper INL (Figure 1 E, F) contained both PV and Prox1 labelled somata. Among them, the double labelled AII amacrine cells were the most numerous (cell densities 3048±497 mm^−2^). Strongly PV positive perikarya, prominent proximal dendrites and no Prox1 expression identified a population of wide-field amacrine cells (PV-positive wide-field amacrine cells, PV-wfACs). Their mean density was 304±57 mm^−2^ (n = 68 ROIs), about one tenth of the AII amacrines. Occasionally, Prox1-positive nuclei appeared among amacrine cells that lacked PV immunoreactivity.

### Analysis of overlapping retinal neuron density gradients

Most retinal neuron types populate the entire retinal sheet, but their densities, dendritic field sizes and interconnections may vary along certain axes such as the central-peripheral, ventral-dorsal or nasal-temporal axes (Ahnelt and Kolb 2000; Bleckert et al. 2014; Nadal-Nicolás et al. 2020; Camerino et al. 2021). Comparing the densities of certain cell types between experimental groups is therefore complicated by the factor of location within the retina. Even measurements from comparable locations of different retinas are subject to variance due to differences of retinal topography between individual animals (Danias et al. 2002; Ortín-Martínez et al. 2010; Galindo-Romero et al. 2013) as well as tissue distortion and uncertainty of anatomical directions in the specimen. Moreover, the concept of eccentricity is not easy to define in rodents. The distance from the optic nerve head is an inherently poor predictor of cell density because (1) the optic nerve head does not correspond to the center of the visual field, (2) cones and ganglion cells in typical rodent retinas have their maximum density dorsally from the optic nerve head, and (3) isodensity contours are elongated horizontally resembling a visual streak. Indeed, only AII amacrine cells showed weak negative correlation (r = −0.25, p = 0.047) with distance from the optic disk, the other correlations being non-significant. Therefore, we sampled the retinas at various locations, which covered statistically the same range of distances from the optic disk for control and treated animals (Z = 0.754, p = 0.62, Kolmogorov-Smirnov test). In the following, we begin with the analysis of the relationship between the different cell densities.

#### The densities of certain retinal cell types are strongly correlated

Figure 2 shows the pooled density measurements for each ROI sampled from retinas of control and VPA treated animals. It is important to note that these density distributions reflect cell density variations both between animals and within animals (between-ROIs). Furthermore, the density variations of the cell types do not seem to be independent but suggest complex associations between them. At the current stage of the analysis, we will describe these relationships, whereas treatment effects, between-animal and within-animal associations will be considered in the next section.

**Figure 2.**
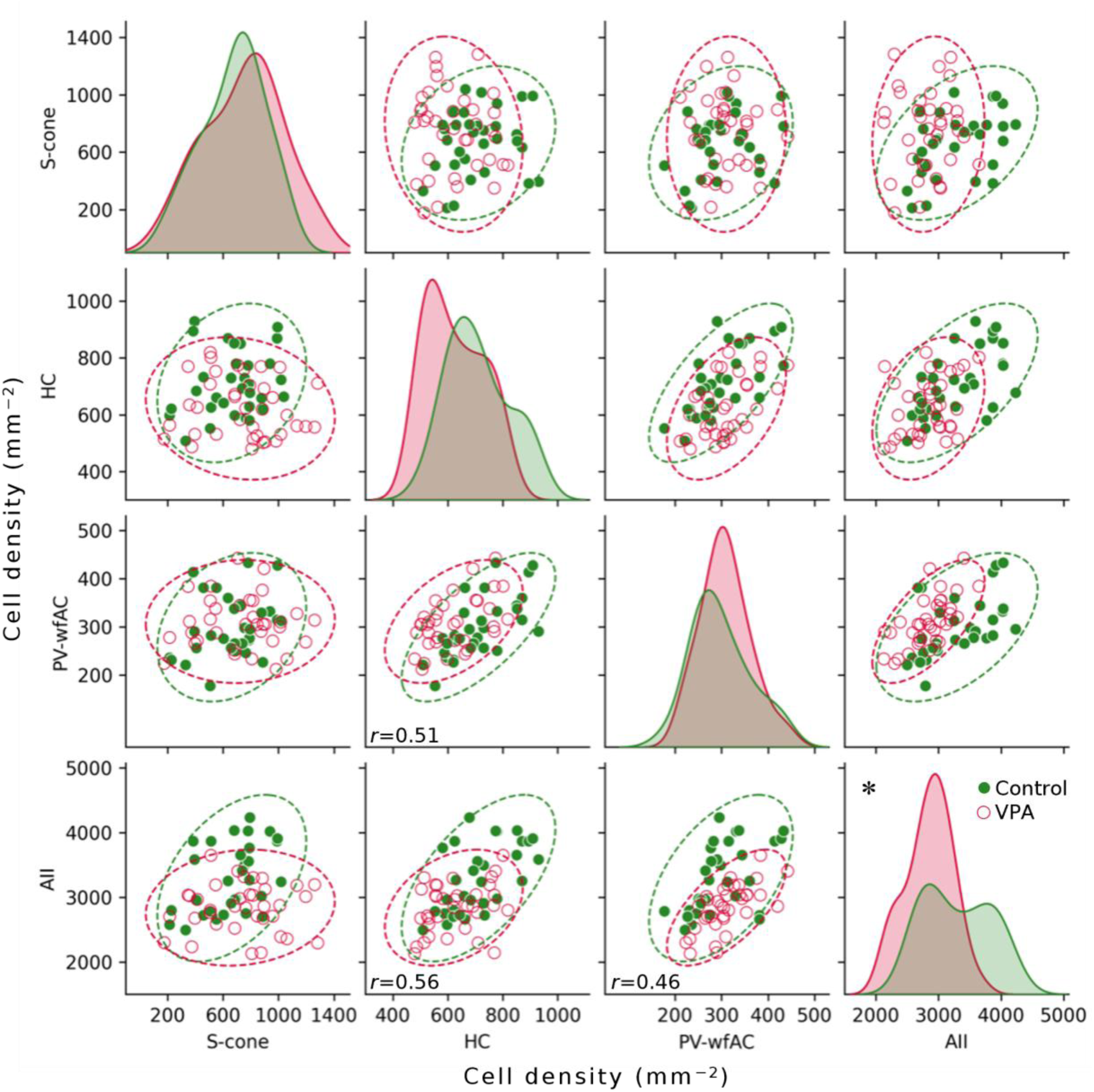
Densities of S-cones, horizontal cells (HC), parvalbumin-positive wide-field (PV-wfAC) and AII amacrine cells are shown in mm^−2^ units. In the scatter plots, each data point corresponds to a ROI either from VPA treated (open red circles) or control (filled green circles) rats. Overall, the cell densities of HCs and amacrine cells were mutually correlated, while S-cones were independent (Pearson’s r values of significant correlations of the pooled data set are shown, Table 3). Distributions of cell densities are shown on the diagonal as kernel density estimates. * Mean AII amacrine cell densities were significantly lower for VPA model rats (LME model, p<0.05, Table 2). Ellipses (red and green for VPA-treated and control animals, respectively) are two-dimensional projections of the 95% confidence intervals for the multivariate mean vectors.

**Table 2.**
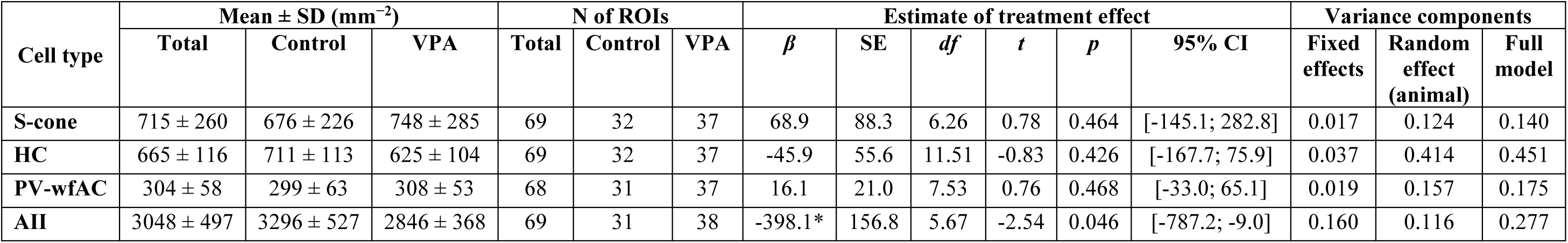
Comparison of mean cell densities. The coefficient β represents the estimated difference between the groups (VPA − Control) from a linear mixed-effects model (LME) with animal identity as a random effect. The numbers of animals were 4 and 6 for the Control and VPA groups, respectively. CI, confidence interval for β; * p<0.05.

| Cell type | Mean ± SD (mm<sup>-2</sup>) | | | N of ROIs | | | Estimate of treatment effect | | | | | | Variance components | | |
| --- | --- | --- | --- | --- | --- | --- | --- | --- | --- | --- | --- | --- | --- | --- | --- |
| | Total | Control | VPA | Total | Control | VPA | $\beta$ | SE | df | t | p | 95% CI | Fixed effects | Random effect (animal) | Full model |
| S-cone | 715 ± 260 | 676 ± 226 | 748 ± 285 | 69 | 32 | 37 | 68.9 | 88.3 | 6.26 | 0.78 | 0.464 | [-145.1; 282.8] | 0.017 | 0.124 | 0.140 |
| HC | 665 ± 116 | 711 ± 113 | 625 ± 104 | 69 | 32 | 37 | -45.9 | 55.6 | 11.51 | -0.83 | 0.426 | [-167.7; 75.9] | 0.037 | 0.414 | 0.451 |
| PV-wfAC | 304 ± 58 | 299 ± 63 | 308 ± 53 | 68 | 31 | 37 | 16.1 | 21.0 | 7.53 | 0.76 | 0.468 | [-33.0; 65.1] | 0.019 | 0.157 | 0.175 |
| AII | 3048 ± 497 | 3296 ± 527 | 2846 ± 368 | 69 | 31 | 38 | -398.1* | 156.8 | 5.67 | -2.54 | 0.046 | [-787.2; -9.0] | 0.160 | 0.116 | 0.277 |

**Table 3.** Correlation statistics of cell densities in the retinas of control and VPA treated rats, as well as in their pooled data. Each value in the table corresponds to a panel in Figure 2. In the blocks Total, Control and VPA, the Pearson’s correlation coefficient (r) and sample size (n) are shown. * p<0.05, ** p<0.001. For Fisher Z-test, p-values and the underlying Z-values are shown.

| Cell type | Pearson's correlation |  |  |  |  |  |  |  |  |  |  |  | Fisher Z-test<br>VPA vs. Control |  |  |  |  |  |
| --- | --- | --- | --- | --- | --- | --- | --- | --- | --- | --- | --- | --- | --- | --- | --- | --- | --- | --- |
|  |  | Total |  |  |  | Control |  |  |  | VPA |  |  |  |  |  |  |  |  |
|  |  | S-cone | HC | PV-wfAC | AII | S-cone | HC | PV-wfAC | AII | S-cone | HC | PV-wfAC | AII |  | S-cone | HC | PV-wfAC | AII |
| S-cone | r | 1 |  |  |  | 1 |  |  |  | 1 |  |  |  | p |  |  |  |  |
| HC |  | -0.025 | 1 |  |  | 0.200 | 1 |  |  | -0.097 | 1 |  |  |  | 0.235 |  |  |  |
| PV-wfAC |  | 0.125 | 0.512** | 1 |  | 0.270 | 0.679** | 1 |  | -0.007 | 0.498* | 1 |  |  | 0.273 | 0.278 |  |  |
| AII |  | 0.121 | 0.556** | 0.460** | 1 | 0.444* | 0.592** | 0.523* | 1 | 0.024 | 0.315 | 0.660** | 1 |  | 0.078 | 0.167 | 0.409 |  |
| S-cone | n | 69 |  |  |  | 32 |  |  |  | 37 |  |  |  | Z |  |  |  |  |
| HC |  | 69 | 69 |  |  | 32 | 32 |  |  | 37 | 37 |  |  |  | 1.187 |  |  |  |
| PV-wfAC |  | 66 | 66 | 68 |  | 31 | 31 | 31 |  | 35 | 35 | 37 |  |  | 1.097 | 1.084 |  |  |
| AII |  | 67 | 67 | 67 | 69 | 31 | 31 | 31 | 31 | 36 | 36 | 36 | 38 |  | 1.764 | 1.380 | −0.826 |  |

Bivariate correlations of cell densities are shown in Table 3 (“Total” correlation matrix). S-cone densities were not significantly correlated with any of the other cell types. On the other hand, correlations were significant between horizontal cells and the two amacrine cell types, a pattern indicating a common principle (a latent variable) shaping their distributions.

The correlation structures between cell densities were similar when the data were split into control and treated groups, both groups showing S-cone densities uncorrelated with the other cell types (see ‘Control’ and ‘VPA’ correlation matrices in Table 3. Correlation matrices as well as covariance matrices for treated animals were statistically similar to those of the controls as confirmed by Jennrich’s test (*χ^2^* = 8.076, *p* = 0.233) (Jennrich 1970; Vermorken 2008) and Box’s M-test (M = 15.32, *p* = 0.161), respectively. Fisher’s Z-test revealed no significant differences between any individual pairs of correlation coefficients from control and treated retinas (Fisher *Z*-test in Table 3). Altogether, these results suggest that the VPA treatment did not alter the their mutual relationships of cell densities across the sampled ROIs. It is important to note that the similarity was in their relationships but not necessarily in the actual densities, which we will consider below.

Next, we used principal component analysis (PCA) of the multivariate dataset, where each ROI was characterized by four density values. PCA aims to express the original data in terms of new variables (principal components) that are uncorrelated (orthogonal to each other) and capture the maximum variance in the data. The first two principal components (PC1 and PC2) captured 73% of variance (Table 4). PC1 had equally high loadings for the densities of horizontal cells and the two amacrine types suggesting similar density distributions across animals and retinas. Scores on the PC1 axis (Figure 3) thus quantified the latent variable shaping the common variation of these three cell types in our dataset. The second principal component (PC2) on the other hand was almost exclusively associated with S-cone density. This suggested an independent distribution pattern of S-cones from the other cell types.

**Figure 3.**
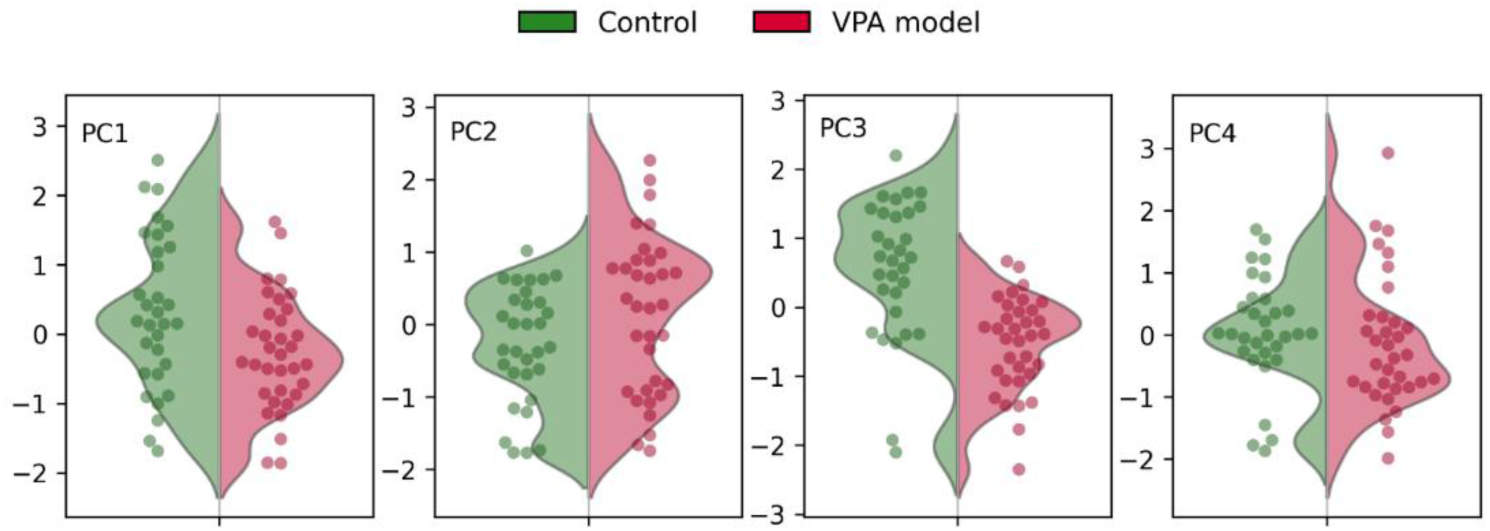
Distributions of principal component (PC) scores of ROIs compared between control (green) and VPA treated (red) rats represented by kernel density estimates. Individual ROIs shown by data points are jittered to avoid overlap. Linear mixed-effects modeling revealed a significant difference between the treatment groups along PC3 (p=0.0073) but not along the other PCs.

**Table 4.** Principal component analysis of the densities of four cell types. Control and VPA treated cases were pooled. Coloring of cells illustrates component loading on a scale from blue (−1) through white (0) to red (+1). Note the separation of S-cones into PC2 from the other cell types in PC1.

| Measured densities | Component loadings |  |  |  |
| --- | --- | --- | --- | --- |
|  | PC1 | PC2 | PC3 | PC4 |
| S-cone | 0.172 | 0.976 | 0.042 | 0.131 |
| HC | 0.832 | −0.242 | 0.093 | 0.490 |
| PV-wfAC | 0.798 | 0.036 | −0.573 | −0.184 |
| AII | 0.822 | 0.006 | 0.453 | −0.345 |
| % of total variance explained | 51 | 25 | 14 | 10 |

Interestingly, PC3 - showing moderately high but opposite loadings for PV-wfAC and AII densities - appeared to strongly separate control and VPA model animals (Figure 3), even though PCA was performed without regard to the experimental group. This suggested that the treatment affected a latent component of the amacrine cell distributions that was independent of the global density gradients represented in PC1 and PC2.

#### PCA of retinal cell densities reveals a functionally relevant measure of centrality

The density of all four cell types showed about twofold or greater variation across the data-set. Retinal location is an obvious source of this variance due to the non-uniform topography of most cell types. Any treatment effect is superimposed onto this natural variation, and to detect it, the effect of location should be subtracted. Since the distance from the optic disk proved to be a poor predictor of cell densities (see above) and rigorous registering retinas from different animals at the microscopic scale is fraught with difficulties of finding appropriate anatomical landmarks, distortions due to flattening the tissue and inter-individual variability of retinal topography, we sought a more reliable location measure.

A functionally relevant measure of retinal location is the density of major cell classes with a center-periphery gradient including photoreceptors and ganglion cells; a similar topography has been found for AII amacrine cells in the rat (Wässle et al. 1993; Gábriel et al. 2004; Liu et al. 2022). We argue that the first principal component (PC1) of cell densities obtained above can be reasonably interpreted as a latent factor of ‘centrality’ because it captures the correlated density variations of HCs and AII amacrines, both of which are constituents of most retinal pathways. Thus, for each ROI, the score on PC1 offers a measure of proximity to the shared density peak(s) of multiple cell types (with very small contribution from S-cones, Table 4).

Further support for PC1 as a useful location measure was gained when plotting its value onto the retinal coordinates of ROIs. When ROIs of all animals were aggregated, a region of high PC1 scores was observed in the dorsal retina, which broadly corresponded to the region of highest densities for HCs, and both amacrine cell types (Supplementary Figure S2). Thus, the analysis of complex topographical relationships between multiple retinas could be simplified to the consideration of a simple, one-dimensional variable.

### Distinctive features of retinal cell density in VPA treated animals

#### VPA treatment selectively affects retinal amacrine cell organization

Figure 2 illustrates several features that distinguish VPA-treated animals from controls. As a first approximation, the mean densities of each cell type were compared between the treatment groups by linear mixed-effects (LME) modeling (Eq. 1, Table 2). Here, AII amacrines were reduced by VPA treatment (statistically on the margin of significance), while S-cone, HC and PV-wfAC densities were not significantly different.

The treatment groups were also compared along the principal component axes by using the same LME model. This revealed a significant difference in PC3 scores (p=0.0073) but not along the other PCs (Figure 3). This meant that (1) the global cell densities captured by PC1 and PC2 were not different between the experimental groups, and (2) PC3 captured a treatment effect that was independent of the global density gradients.

Thus, the above analyses revealed two features that differentiated the retinas of VPA-treated offspring from controls: AII amacrine cell density (lower in the VPA group) and PC3 (reflecting opposite density changes in the two amacrine cell types). None of the above analyses did, however, formally adjust for sampling potentially non-corresponding locations in the retinas.

#### Centrality-adjusted analysis reveals opposite treatment effects in the two amacrine cell types

To compare equivalent locations in the retinas, we built an LME model that assumed linear relationship between centrality (PC1) and cell density in addition to the treatment effect. Since PC1 was derived from the pooled data of all animals, it may reflect correlations within animals (i.e. retinas) as well as between animals. To separate the effects of between-animal and within-animal variation in centrality, PC1 was decomposed into animal-specific means (*PC1_mean_*, between-animal component) and deviations from those means (*PC1_cent_* = *PC1*−*PC1_mean_*, within-animal component), and both terms were entered as covariates. Thus, the final model assumed three fixed effects: treatment, *PC1_mean_* and *PC1_cent_* along with the random variation of mean density between animals (Eq. 2). This model was fitted separately to each of the four cell densities.

The parameter estimates of the two-covariate LME model are shown in Table 5. None of the predictors had a significant effect on S-cone density, therefore in the following, we concentrate on the results for the remaining cell types. Also, the model fits were remarkably good with R^2^ (conditional) values over 0.75 for HCs and amacrine cells while the fit to S-cone densities was rather poor (Table 6). Most of this explained variance concentrated in the fixed effects indicating that they gave a good description of the data at the level of ROIs.

**Table 5.** Results of fitting a linear mixed-effects model accounting for the joint effects of centrality (PC1) and prenatal VPA treatment on retinal cell densities, with animal identity as a random effect (Eq. 2). The centrality was partitioned into between-animal and within-animal variation using each animal’s mean PC1 (PC1_mean_) and animal-centered PC1 (PC1_cent_) scores as covariates, respectively. The coefficients (β) for Treatment represent the estimated differences between the groups (VPA − Control) after adjustment for centrality. SE, standard error of β; df, degrees of freedom with Satterthwaite’s correction; CI, confidence interval for β; * p<0.05, ** p<0.001.

| Cell type | Fixed effect | Fixed-effect estimates |  |  |  |  |  |
| --- | --- | --- | --- | --- | --- | --- | --- |
| | | $\beta$ | SE | df | $t$ | $p$ | 95% CI |
| S-cone | Treatment | 84.0 | 96.3 | 8.62 | 0.87 | 0.406 | [-135.3; 303.3] |
| | $PC1_{mean}$ | 69.3 | 81.5 | 8.95 | 0.85 | 0.418 | [-115.3; 253.9] |
| | $PC1_{cent}$ | 68.4 | 35.3 | 55.92 | 1.94 | 0.058 | [-2.4; 139.2] |
| HC | Treatment | -3.6 | 33.1 | 6.43 | -0.11 | 0.918 | [-83.4; 76.3] |
| | $PC1_{mean}$ | 129.6* | 27.9 | 6.49 | 4.64 | 0.003 | [62.5; 196.7] |
| | $PC1_{cent}$ | 78.9** | 7.7 | 53.71 | 10.24 | $3.13 \times 10^{-14}$ | [63.4; 94.3] |
| PV-wfAC | Treatment | 35.2* | 11.8 | 7.96 | 2.97 | 0.018 | [7.9; 62.5] |
| | $PC1_{mean}$ | 42.0* | 10.0 | 8.23 | 4.20 | 0.003 | [19.0; 65.0] |
| | $PC1_{cent}$ | 52.8** | 4.2 | 55.43 | 12.63 | $5.91 \times 10^{-18}$ | [44.4; 61.1] |
| AII | Treatment | -288.0* | 114.3 | 8.29 | -2.52 | 0.035 | [-550.0; -26.0] |
| | $PC1_{mean}$ | 305.4* | 96.5 | 8.44 | 3.16 | 0.012 | [84.8; 525.9] |
| | $PC1_{cent}$ | 416.4** | 33.8 | 55.51 | 12.31 | $1.69 \times 10^{-17}$ | [348.6; 484.2] |

**Table 6.** Estimates of the proportions of observed total variance explained by the best-fit linear mixed-effects model for each cell type (Table 5). Marginal R^2^denotes the proportion of total variance explained by the fixed effects. Conditional intraclass correlation coefficient (ICC) denotes the proportion of total variance attributable to the random effect (animal identity) after accounting for the fixed effects. Conditional R^2^ denotes the proportion of variance explained by the full model, including both fixed and random effects.

| Cell type | Variance components |  |  |
| --- | --- | --- | --- |
| | Fixed effects (marginal $R^2$) | Random effect (conditional ICC) | Full model (conditional $R^2$) |
| S-cone | 0.07 | 0.14 | 0.21 |
| HC | 0.69 | 0.13 | 0.82 |
| PV-wfAC | 0.72 | 0.05 | 0.77 |
| AII | 0.74 | 0.07 | 0.81 |

The effects of between-animal and within-animal variation in centrality were both significant, suggesting that the mean densities of the same cell types remained correlated despite the variation of mean centrality between animals. The p-values for between-animal variation were greater by several orders of magnitude likely owing to the smaller effective sample size for estimating its effect.

The treatment-related density differences estimated by the model can be visually summarized as two parallel regression lines corresponding to the control and treated groups, respectively, with their vertical distance showing the estimated difference in mean cell density at comparable locations of control and treated retinas (Figure 4). The analysis showed that these differences were statistically significant for both amacrine cell types, but the direction of change was opposite (Table 5). For PV-wfACs, the change in density due to prenatal VPA treatment was positive, representing an 11.6% increase relative to the mean (Table 2). Conversely, AII amacrines were decreased by 9.4% compared to controls. In the same analysis, the VPA effect was non-significant for HCs or S-cones (Table 5, Figure 4).

**Figure 4.**
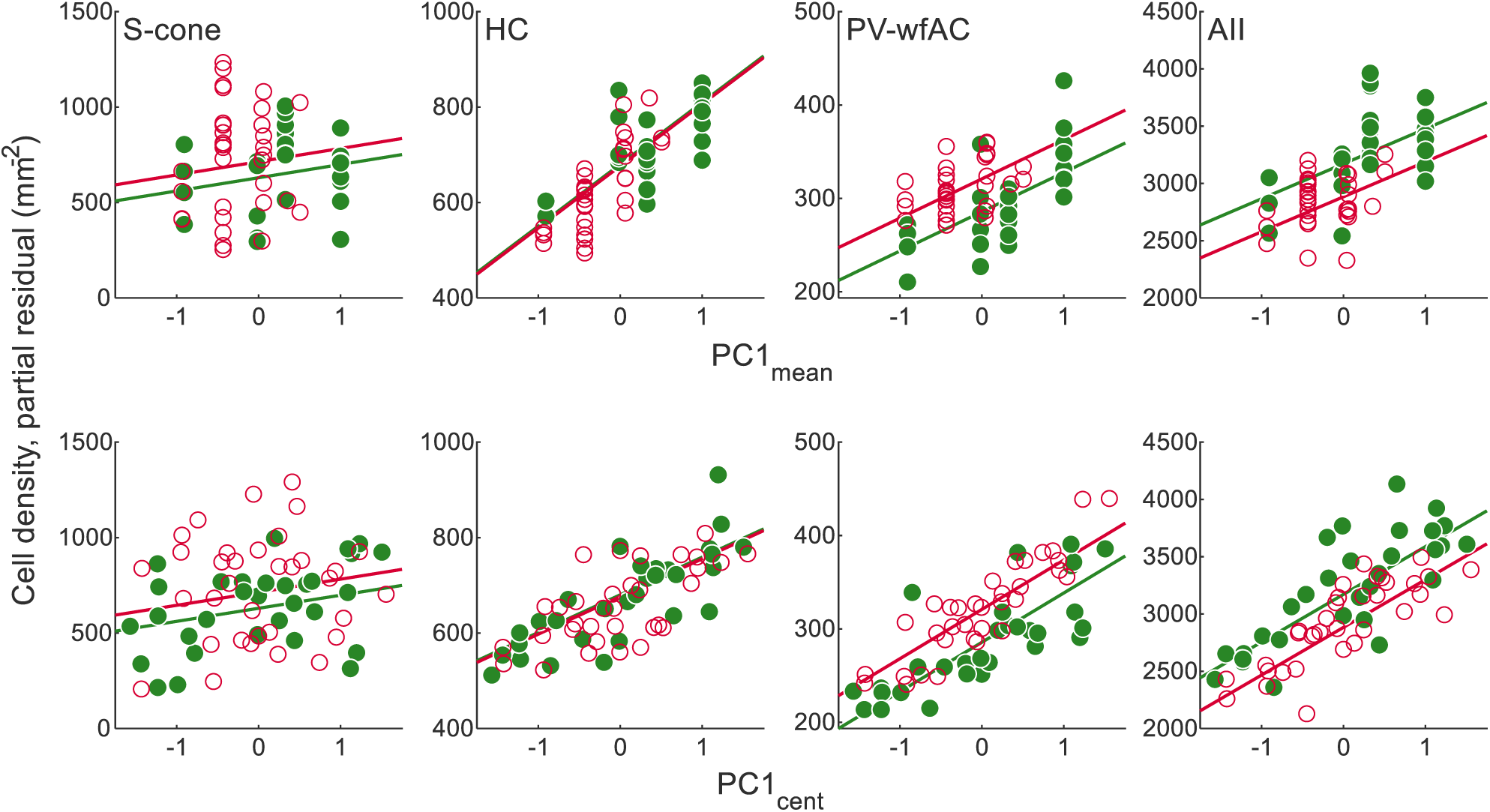
Partial residual plots of cell densities of S-cones, HC, PV-wfAC and AII amacrines plotted as a function of centrality (PC1), used here as a measure of location derived from global cell density. Centrality was decomposed into between-animal (PC1_mean_) and within-animal variation (PC1_cent_) and entered as fixed effects in the linear mixed-effects model (Eq. 2, Table 5). In the top row, the effect of PC1_mean_ is shown while holding PC1_cent_ at zero; in the bottom row, the effect of PC1_cent_ is shown while holding PC1_mean_ at zero. Each symbol represents a retinal ROI from a control (filled green circles) or VPA treated (open red circles) rat. The plotted ordinate is the component-plus-residual value, so the vertical distance of each point to the fitted regression line equals the model residual. Regression lines for the control (green) and VPA (red) groups are derived from the model. Thus, the vertical distance between the lines corresponds to the estimated treatment effect, which was significant for PV-wfACs and AII amacrines.

### Mosaic regularity is unaffected in the VPA-model

The individual cells of each neuronal cell type are arranged relative to each other in a more or less regular cell mosaic. These mosaics are never geometrically perfect: the lateral distances between neighboring cells vary, with larger variance pointing to a less regular mosaic. We quantified cell mosaic regularity for each ROI and cell type by the regularity index derived either from nearest-neighbor distances (NNRI) or from the areas of Voronoi-domains of each cell (VDRI). Regularity indices (RI) are calculated as the mean divided by the standard deviation of the respective measure so that more regular cell arrangement and thus, more uniform nearest-neighbor distances or Voronoi-domain areas result in a higher index. The key difference between the two indices is that VDRI accounts for distances to all neighboring cells, while NNRI only considers the nearest neighbor.

Taking all retina samples together, the mosaics of each cell type had characteristic regularities with their median regularity indices increasing in the order S-cones < PV-wfAC < AII amacrines < horizontal cells (Figure 5). Kruskal-Wallis test showed significant main effect of cell type for both NNRI and VDRI (p<0.0001). Subsequent pair-wise comparison revealed significant differences except for the NNRIs of AII amacrines and HCs (*p*>0.05 with Bonferroni correction). A weak positive correlation was observed between the density of AII amacrines and their regularity (Spearman’s *ρ* = 0.26 for NNRI and 0.28 for VDRI, p<0.05). This phenomenon may be due to the known effect of crowding, whereby a higher density cell mosaic is forced to a more regular pattern (Reese and Keeley 2015).

**Figure 5.**
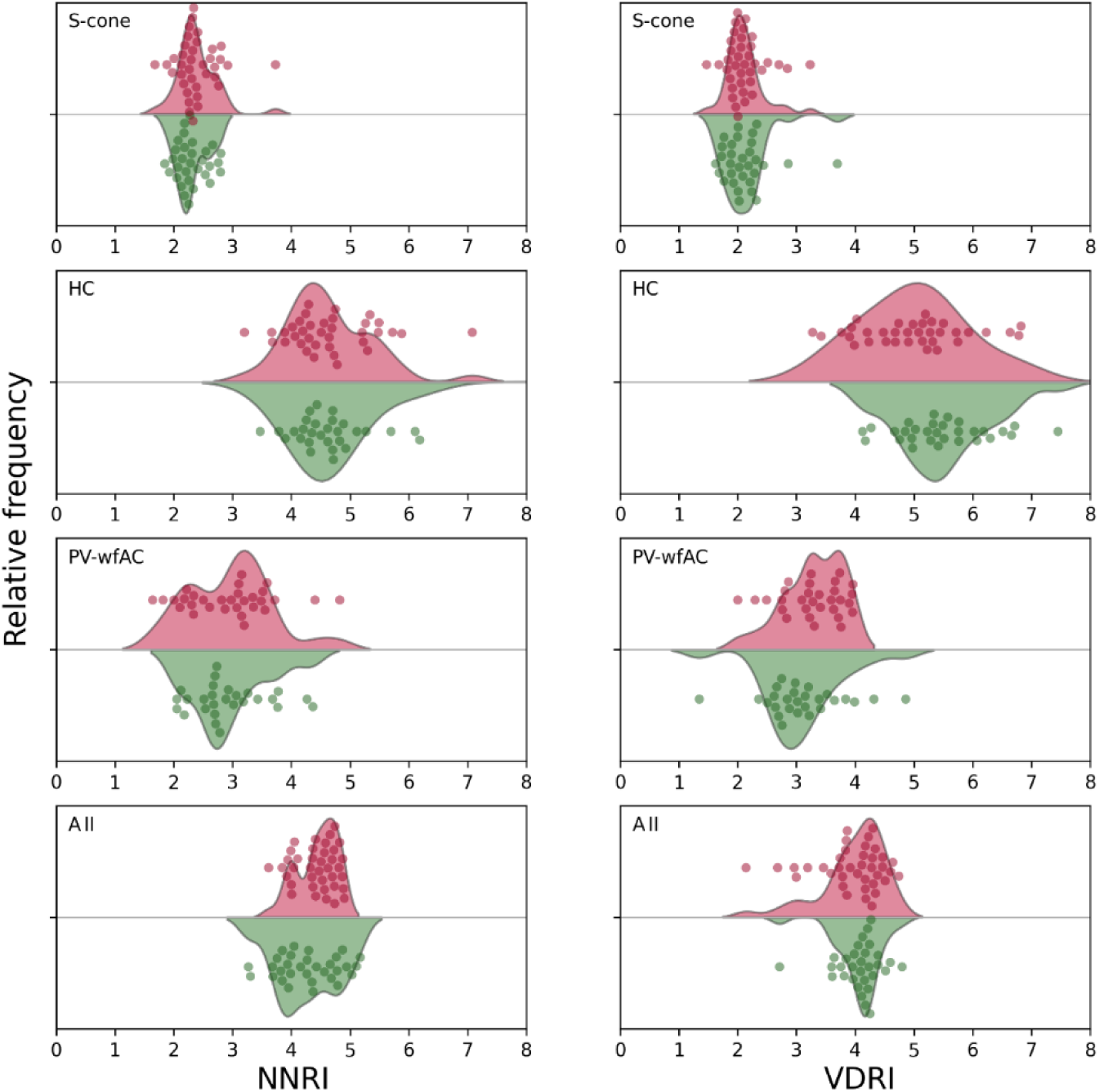
Distribution of regularity indices calculated from nearest neighbor distances (NNRI) or Voronoi domain areas (VDRI) for each cell type and region of interest (ROI) represented by kernel density estimates. Individual ROIs shown by data points are jittered to avoid overlap. No significant differences of the mean regularity indices between VPA-treated (red) and control (green) animals were found.

Mosaic regularities of the control and VPA groups were compared by fitting a LME model for each cell type, with treatment as the fixed effect and animal identity as the random factor. The results were uniform for all cell types: neither NNRIs, nor VDRIs of the treated animals showed a significant difference from controls.

## Discussion

We analyzed the densities and regularity indices of multiple retinal neuron types in the VPA induced rat model of autism spectrum disorder. We developed a statistical model whereby equivalent locations could be compared between multiple retinas by considering a simple, one-dimensional centrality variable. The main finding of the current study is that neuronal cell types of the retina are differentially affected in the VPA model. These changes were predominant among amacrine cells, while horizontal cells and S-cones were not affected. The treatment-related differences could not be explained by a global reduction in cell density because they were opposite in AII amacrine cells (density decrease) and parvalbumin-positive wide-field amacrines (density increase). We found no evidence for major alterations in mosaic regularity.

### Interpretation and limitations of the analysis

The multivariate dataset obtained from the ROIs had the advantage that patterns of density relationships could be identified. Mapping multiple ROIs within retinas not only increased the precision of the animal’s mean cell-density estimates; it also allowed us to study within-animal density variations. Most importantly, it allowed us to adjust our estimates of the treatment effect for the most important confounding factor encountered when comparing different retinas, namely location. Mapping multiple locations did not, however, compensate for the relatively small number of animals. Replication in a larger cohort will therefore be important to establish the robustness of the findings and obtain more precise estimates of effect size.

The concurrent mapping of multiple cell types at multiple retinal locations enabled inferences based on the relationships among their densities that would not have been possible from separate analyses of individual cell populations. However, the scope of these inferences is constrained by the choice of cell types included in the study. The populations examined here were selected to represent different retinal processing systems and levels of circuitry (see Introduction). Future studies incorporating additional cell types, including other photoreceptor classes and ganglion cells, may uncover more general organizational principles and permit testing of hypotheses generated by earlier work (Guimarães-Souza et al. 2019).

While we believe that calling the first principal component of the multivariate cell density space “centrality” is well justified, factors other than location might have contributed to it. One potential contributor would be any between-animal variation of cell density, whether due to random variation, sampling non-corresponding locations or treatment-related differences. Differential tissue distortion due to shrinkage or flattening could also cause differences in global cell density between ROIs even if cell distributions were completely flat. Importantly however, PC1 would remain a useful covariate to adjust in such cases. Furthermore, if treatment effects were captured by PC1, it would have been identified by the comparison of mean PC1 scores, as they were revealed for PC3 (Figure 3).

Nevertheless, one can envisage treatment effects that would remain undetected by our analysis. These include alterations that do not affect the mean cell densities, the derived latent variables, or the relationships among them. At the extreme, even a wholesale rearrangement of retinal topography could escape detection if the measured density relationships were preserved.

### Relationship to previous studies of cell mosaics

Our measurements of cell densities (Table 2) broadly agree with previous studies (see Wässle *et al*., 1993; Gábriel *et al*., 2004; Liu *et al*., 2022 for AII and PV-positive wide-field amacrines, Chu *et al*., 1993; Peichl & González-Soriano, 1994 for HCs and Arango-Gonzalez et al. 2010; Ortín-Martínez et al. 2010 for S-cones, Table 7). However, the concurrent labelling of these cell types and the analysis of their density covariation allowed some insight into their topographic relationships.

**Table 7.**
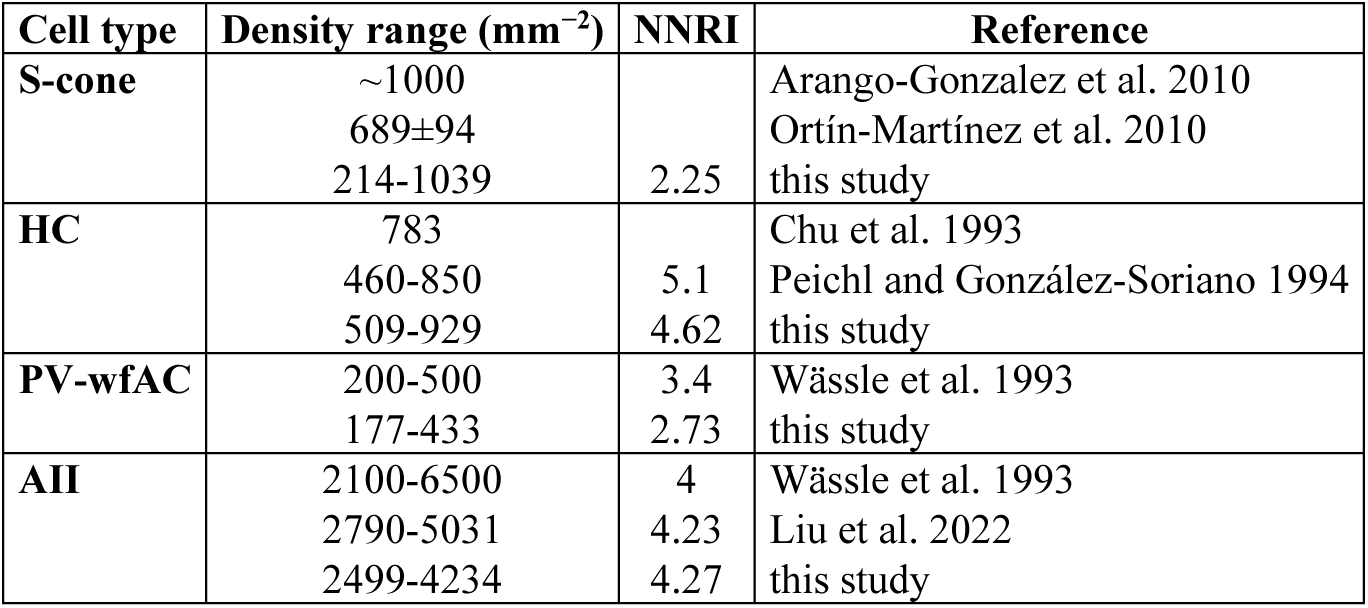
Comparison of cell densities and regularity indices reported in various studies. Density values from the current study represent control animals. Density values from other studies are either means of the reported values or mean ± standard deviation. NNRI values are shown as medians for the current study and as means for previous studies.

First, our data corroborate results from several species that S-cones show a unique topography, which is independent of the distribution of M-cones (Martin and Grünert 1999; Szél et al. 2000; Ahnelt et al. 2006) and their downstream targets involved in spatial vision. In rats, S-cone density gradually decreases from the ventral toward the dorsal retina, and additional hot spots may exist in the extreme periphery (Szél and Röhlich 1992; Ortín-Martínez et al. 2010). This characteristic spatial distribution likely explains why S-cones segregate from the rather dorsally peaking cell types into orthogonal principal components of cell-density covariation.

Second, quantitative estimates of the S-cone mosaic regularity in the laboratory rat have not been published before to our knowledge. Our median NNRI of 2.25 is however, close to figures reported for another murine rodent, the Nile grass rat (between 1.84 and 2.03, Gaillard et al. 2009). In comparison to M/L-cones, the regularity of the S-cone mosaic was significantly lower in the same study. This is consistent with our current finding that the S-cone mosaic had the lowest regularity among the four cell types. Furthermore, this irregularity seemed to be consistent across the retina as suggested by the narrow spread of regularity indices (Figure 5).

Third, the strong covariation of PV-positive wide-field amacrines with AII amacrine density in our data suggests that they also follow a center-periphery gradient. Topographical data for PV-wfACs has not been available so far (Wässle et al. 1993; Liu et al. 2022).

Finally, the correlation of horizontal cells with AII densities is difficult to reconcile with earlier studies of HC topography in the rat (Chu et al. 1993; Peichl and González-Soriano 1994) that found an apparent lack of a clear center-periphery gradient. Guinea pigs in contrast, have HC density peaks in the superior retina, gerbils in the inferior retina (Peichl and González-Soriano 1994), while mice possess a complex topography with maxima in the inferior and equatorial regions (Camerino et al. 2021; Spinelli et al. 2024). Altogether, these findings suggest that the topography of horizontal cells is adaptable under various phylogenetic or ontogenetic selection constraints.

### Developmental insights

The various groups of retinal neurons proliferate and differentiate in a sequential manner during prenatal and early postnatal development (Rapaport et al. 2004; Hussey et al. 2022). Although we performed our analysis on adult animals, it is worth considering hypothetical developmental scenarios that could potentially account for the observed alterations.

For the rat autism model, VPA is given during the first wave of retinal neurogenesis, during which ganglion cells, horizontal cells and cone photoreceptors reach their maximum rate of cell division (Phase 1 of Rapaport *et al*., 2004). In proliferating neural progenitors, VPA induces arrest in the G1-phase of the cell cycle and reduces mitotic activity, while simultaneously promoting neuronal differentiation (Hall et al. 2002; Jung et al. 2008). These effects have been demonstrated in cortical and hippocampal progenitors and could potentially extend to retinal neuroblasts, which share similar regulatory programs. An early mitotic cycle exit of the cell types proliferating during Phase 1 could therefore be reflected in a reduced number of S-cones and horizontal cells. In our experiments however, the major change occurred in late-born amacrine cells while HC numbers were only marginally decreased, and S-cone numbers were not affected. This pattern is consistent with the possibility that VPA’s developmental impact may extend beyond early proliferative events to later differentiation stages.

The selective reduction of AII amacrine cells observed here may reflect epigenetic reprogramming of glycinergic amacrine precursors or altered signaling environments that compromise survival during Phase 2 of retinal neurogenesis (Rapaport et al. 2004) or may bias lineage commitment during differentiation. In contrast, the relative increase in PV-wfAC numbers could result from a competitive advantage within the amacrine cell population under disrupted developmental conditions or represent a compensatory response to maintain inhibitory balance.

Finally, the absence of detectable changes in mosaic regularity despite significant alterations in cell density is consistent with the possibility that the mechanisms governing spatial patterning are relatively resistant to the developmental effects of VPA. Homotypic interactions that are mediated by contact-dependent repulsion and dendritic field exclusion, are thought to operate after the major phases of proliferation and differentiation, during a postmitotic period of mosaic refinement (Raven and Reese 2002; Eglen and Galli-Resta 2006; Reese 2011; Kay et al. 2012). For many retinal cell types, including horizontal and starburst amacrine cells, this refinement occurs well after cell cycle exit, often during the first postnatal week in rodents, when tangential dispersion and dendritic tiling adjust local spacing (Raven and Reese 2003; Kozlowski et al. 2024). These self-organizing processes act locally and iteratively, enabling each cell type to maintain characteristic neighborhood relationships even when the absolute number of cells is perturbed. Thus, one possible interpretation is that developmental processes operating after cell production may have compensated for earlier perturbations, ultimately contributing to preserved regularity in adulthood.

Morphological differences observed in the adult retina may reflect an altered morphogenetic environment within the developing central nervous system, and the atypical retina may arise independently of the atypical behavioral phenotype (Laszlo et al. 2025). However, sensory organs are also active participants in neural development. Before the onset of vision, spontaneous retinal activity contributes to the maturation of visual pathways, whereas after eye opening the retina provides the structured sensory input required for the experience-dependent refinement of cortical circuits. Consequently, retinal abnormalities may not merely be a correlate of atypical brain development, but could potentially contribute to the emergence of altered neural circuitry and behavior. Longitudinal studies tracking ocular development alongside behavioral maturation will be required to distinguish between these possibilities.

### Retinal neuron phenotypes in ASD and its animal models

Evidence from both humans and animal models increasingly supports the notion that the retina is affected in ASD. In human ASD, optical coherence tomography (OCT) studies have reported structural changes that may include both thinning (Emberti Gialloreti et al. 2014) and thickening of the peripapillary nerve fiber layer (García-Medina et al. 2017), as well as thinning of the macular region (Friedel et al. 2024) compared to neurotypical controls. Across these studies, individuals with greater functional impairment (lower intelligence quotient or higher social impairment scores) generally exhibited thinning of the affected retinal layers, although the specific measures of deficit and the strength of the correlations varied. Altogether, the OCT findings point towards a relative decrease of retinal ganglion cell numbers with increased symptom severity. Compatible with these results, a deep learning-based classification study of retinal photographs highlighted the optic disc area as having the highest predictive value for ASD (Kim et al. 2023).

Functionally, a series of studies employing electroretinography (ERG) has provided evidence for reduced b-wave amplitudes in human ASD (Ritvo et al. 1988; Constable et al. 2016, 2020; Lee et al. 2022) implicating inner retinal contributions, particularly within the ON pathway. Others found no significant differences in ERG parameters in high-functioning adults with ASD (Tebartz van Elst et al. 2015; Friedel et al. 2022), highlighting heterogeneity in both ASD phenotypes and experimental protocols. Collectively, findings in ASD patients suggest that while retinal abnormalities are variable, they occur frequently enough to warrant consideration as biomarkers in ASD.

Animal models have provided more direct evidence of retinal changes associated with ASD-like symptoms. In the prenatal valproate model specifically, Guimarães-Souza et al. (2019) reported a reduction in scotopic a-wave amplitude, suggesting outer retinal involvement. Conversely, the AII amacrine cell loss observed in our current study would favor attenuated b-wave amplitudes and/or altered OPs (Keeley et al. 2023; Joachimsthaler et al. 2025). Biochemical evidence (Guimarães-Souza et al. 2019) points to altered excitatory/inhibitory balance: decreased GABA, glutamate decarboxylase, synapsin-1 and fragile X messenger ribonucleoprotein (FMRP), and increased metabotropic glutamate receptor 5 expression. Similarly, the changes in amacrine cell densities in our results are broadly consistent with altered inhibitory neurotransmission, whereby the glycinergic AII amacrine cells, and putative GABAergic wide-field amacrine cells were affected in opposite ways, suggesting pathway specific impairment and caution against assuming uniform effects on inhibitory circuits.

Genetic mouse models with ASD-like phenotypes differ from each other in the pattern of impairments in various organs making comparisons with each other and with the VPA model difficult. Retinal alterations that have been described often seem to involve the rod photoreceptors, which is evidenced by decreased rhodopsin content (Rossignol et al. 2014; Zhang et al. 2019) and decreased scotopic a-wave amplitudes in the ERG (Rossignol et al. 2014; Zhang et al. 2019; Cheng et al. 2020). Electrophysiological evidence for a decreased function of downstream circuits has also been described (reduced scotopic b-wave (Rossignol et al. 2014; Zhang et al. 2019) or oscillatory potentials (Cheng et al. 2020)). In the Engrailed-2 knockout mouse, a complex pattern of retinal alterations was observed, in which reductions in bipolar cell markers and calbindin-positive horizontal cell numbers were accompanied by increased amacrine cell density and elevated parvalbumin mRNA expression (Zhang et al. 2019). The expression of synaptic markers has been quantified by Western-blot in retinas of Fmr-1 knock-out mice, and both decreased and increased levels have been observed (Rossignol et al. 2014).

Altogether, these observations indicate that both outer and inner retinal circuits may be affected in conditions associated with ASD-like symptoms. As noted in the previous section, the changes in amacrine cell numbers observed in our data together with unaffected S-cone photoreceptor numbers indicate direct developmental effects of VPA on inner retinal circuits. However, evidence from the above ASD model systems raises the possibility that the alterations of inner retinal circuits can also be secondary to photoreceptor loss or impaired photoreceptor function (Strettoi et al. 2002; Lee et al. 2021).

The cell types analyzed in this study represent only a fraction of the retina’s neuronal diversity. Their differential vulnerabilities to VPA exposure suggest that other populations - such as bipolar and ganglion cells - may also be affected in complex ways. Future work should extend the analysis to other cell classes and incorporate functional assessments, including ERG protocols optimized for ON-pathway and inner retinal function (e.g., scotopic b-wave, oscillatory potentials, and photopic hill analysis). Such studies would help bridge the gap between cellular alterations and visual function, providing a more comprehensive understanding of retinal involvement in ASD.

It is, however, important to consider the complex etiology of ASD and the inherent limitations of animal model validity when drawing parallels across these findings. These factors constrain the translatability of results between animal models and humans, and even across subgroups of ASD patients with different genetic or environmental backgrounds.

## Acknowledgements

We are grateful to Erzsébet Korona, Rita Illés, Pálma Fogasi, Paula Hirsch and Attila Rácz for technical assistance. The research was performed in collaboration with the István Ábrahám Nano-Bio-Imaging core facility at the Medical School, University of Pécs, Hungary.

## Abbreviations

ASD: autism spectrum disorder
Eq.: Equation
GABA: gamma amino butyric acid
HC: horizontal cell
ICC: intraclass correlation coefficient
INL: inner nuclear layer
NNRI: nearest neighbor regularity index
OCT: optical coherence tomography
PCA: principal component analysis
PRL: photoreceptor layer
PV-wfACs: parvalbumin-positive wide-field amacrine cell
RI: regularity index
ROI: region of interest
VDRI: Voronoi-domain regularity index
VPA: valproic acid

## Declarations

### Funding

The project was supported by the New National Excellence Program of the Ministry for Innovation and Technology (ÚNKP-21-5-PTE-1333) and the Thematic Excellence Program 2021 Health Sub-programme of the Ministry for Innovation and Technology in Hungary, within the framework of project TKP2021-EGA-16 of the University of Pécs.

### Competing interests

The authors have no competing interests or other interests that might be perceived to influence the results and/or discussion reported in this paper.

### Author Contributions

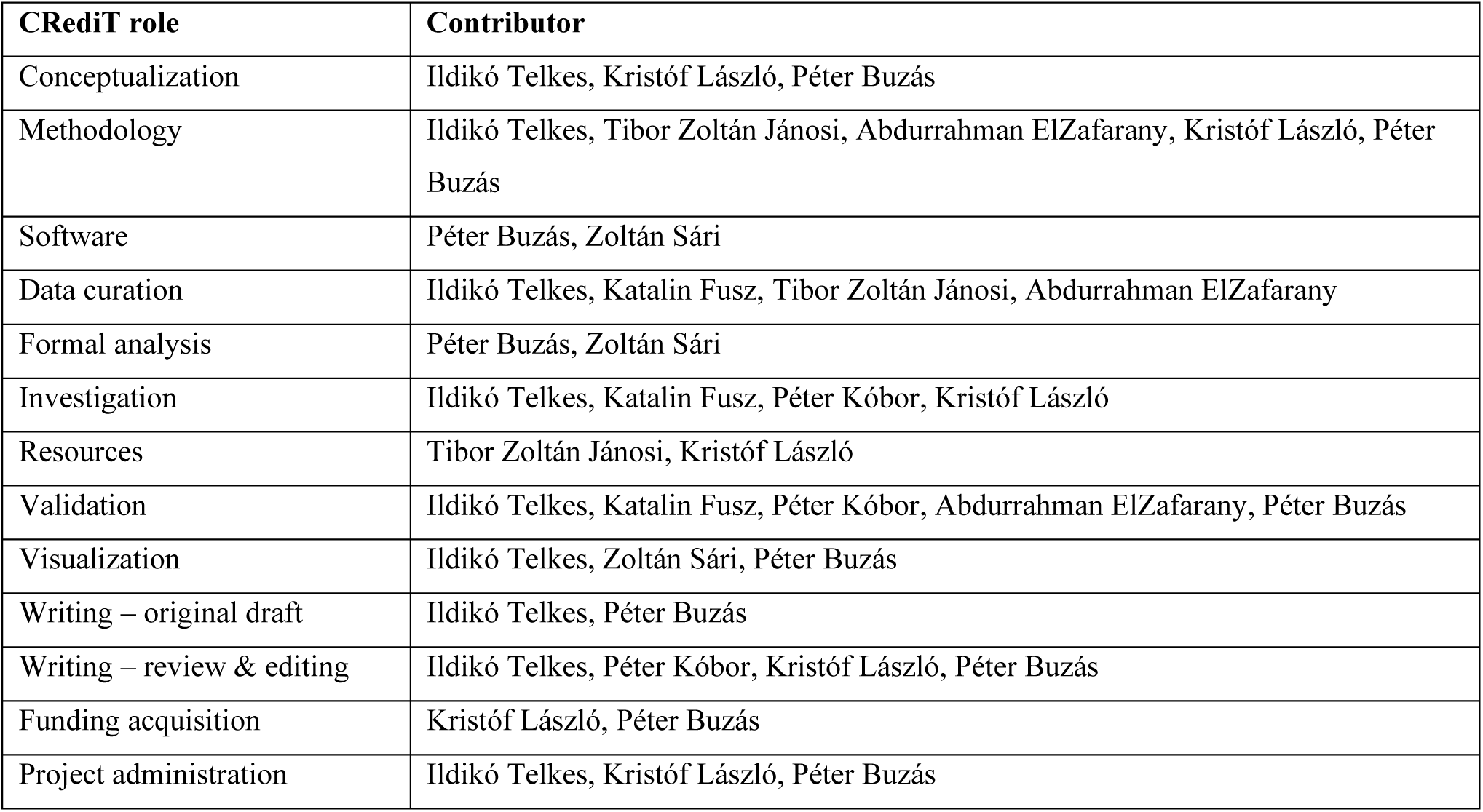

## Notes

### Competing Interest Statement

The authors have declared no competing interest.

### Summary of Updates

Major update of statistical analysis. Substantial changes to Figures 1 and 4. New section on "Interpretation and limitations of the analysis" in the Discussion.

