## Supplementary Figure for "Retinal cell mosaics in the valproate-induced rat model of autism spectrum disorder"

### Supplementary Figures

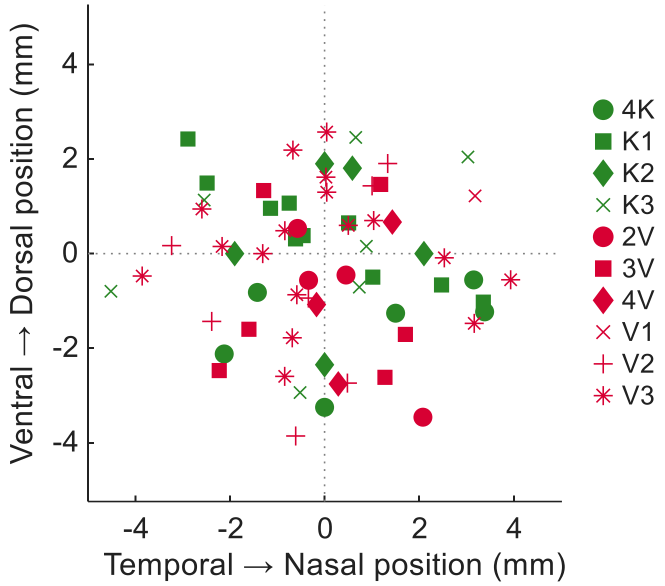


***Supplementary Figure S1.*** *Estimated locations of the sampled regions of interest in anatomical coordinates. The origin represents the optic disc. Markers indicate animal identity. Control animals are shown in green, VPA treated animals are shown in red. Note that retinal locations were not used in the statistical analysis.*

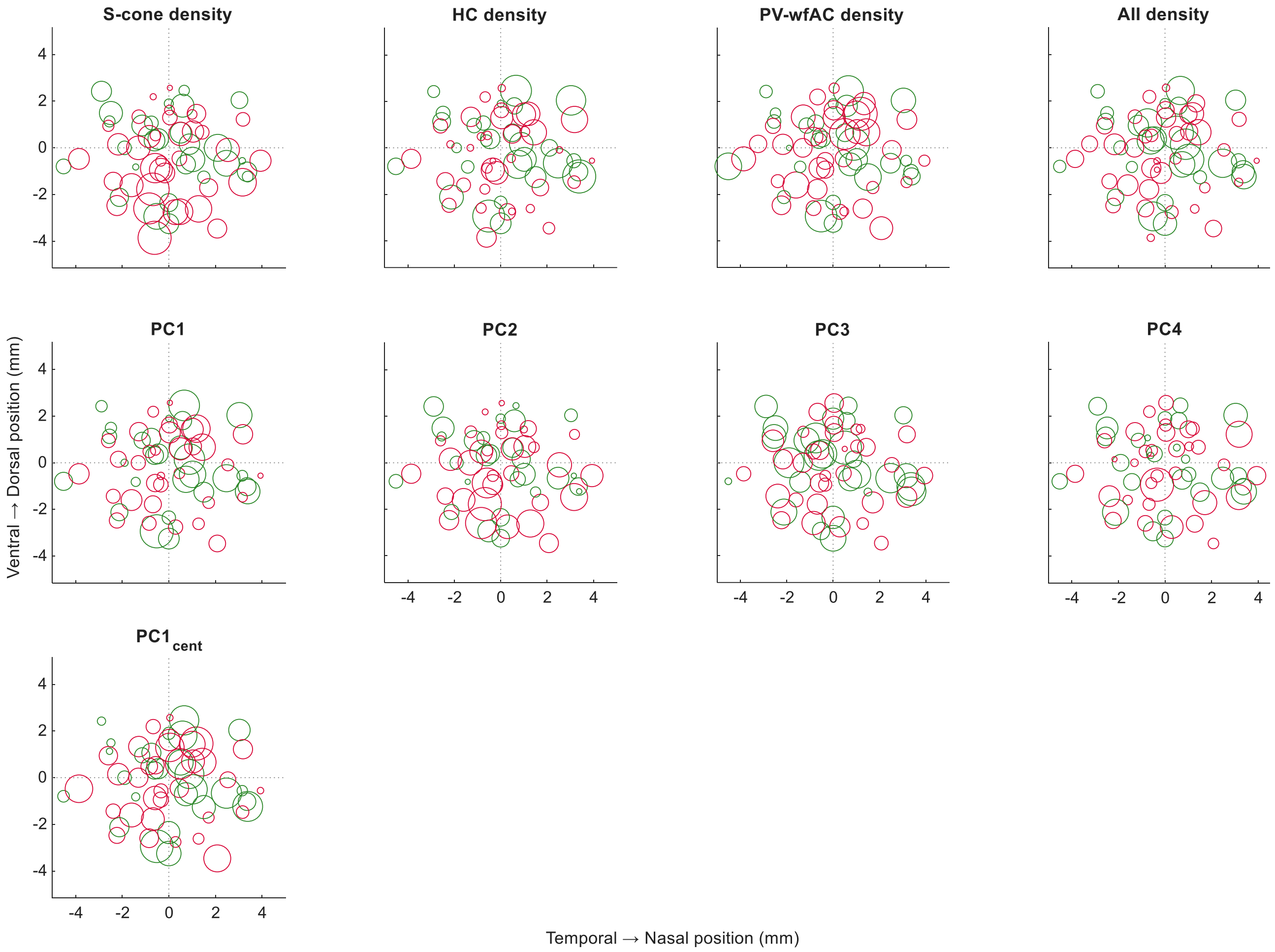


***Supplementary Figure S2.*** *Retinal distributions of the variables shown in the panel titles derived from multiple retinas. The diameter of each circle represents the relative value of the variable, their position corresponds to the estimated retinal coordinates of the corresponding ROI similar to Supplementary Figure S1. The origin represents the optic disc. Data points of control animals are shown in green, VPA treated animals are shown in red.*
